# ProtCell: a foundation model for single-cell proteomics

**DOI:** 10.64898/2026.10.22.760932

**Authors:** Bowen Hu, Zhen Li, Qun Jiang, Qijin Yin, Zeyang Liu, Rui Jiang

## Abstract

Single-cell proteomics provides a direct functional readout of cellular states, but targeted assays measure limited and non-identical protein panels, fragmenting information across experiments. Here, we present ProtCell, a foundation model developed using approximately 380 million cells from 98 datasets for pretraining and evaluation. ProtCell combines protein-identity and continuous-abundance embeddings in a transformer encoder and learns cell- and protein-level representations through masked abundance modeling. ProtCell first connects fragmented profiles through transferable annotation and batch correction while preserving biological organization. It then extends limited profiles through abundance reconstruction and inference of an unmeasured cytotoxic protein program enriched in natural killer cells and memory CD8^+^ T cells. The learned protein dependencies further guide the selection of compact panels that retain information for profile reconstruction and biological classification. ProtCell thus provides a framework for connecting fragmented proteomic profiles, extending information beyond measured panels and guiding more informative measurements.

## Main

Proteins execute most cellular functions and provide a direct readout of signalling, immune recognition and responses to perturbation [1]. Protein abundance and activity cannot be fully inferred from messenger RNA because translation, turnover and post-translational regulation provide additional layers of control [2, 3]. Direct protein measurements are therefore important for resolving the receptors, signalling states and effector molecules that determine how cells interact with and respond to their environment [4, 5]. At single-cell resolution, such measurements have mapped haematopoietic differentiation and drug responses, defined the tumour and immune ecosystems of breast cancer, identified immune states associated with vaccine responsiveness and lupus activity, and revealed aberrant differentiation and surface markers in acute myeloid leukaemia [6–9]. These studies illustrate how single-cell proteomics complements transcript-based atlases by linking cell identity to functional state and clinically relevant phenotypes.

The main limitation of current single-cell proteomic assays is the number of proteins measured in each experiment. Targeted technologies depend on panel design and reagent availability, whereas deeper profiling generally measures fewer cells [4–6, 10, 11]. Panels also differ among studies and platforms, so collectively abundant measurements remain fragmented across partially overlapping protein sets. This fragmentation complicates comparison across experiments, leaves informative proteins unmeasured in individual datasets and makes the choice of markers consequential for future assays. Existing methods support clustering, batch correction and abundance recovery [12–14]. The broader opportunity is to reuse information distributed across panels, so that measurements from one study can inform the interpretation of another. Pretraining on heterogeneous single-cell atlases has supported transferable representations in other molecular modalities [15,16]. For protein measurements, a shared representation could connect cell states across experiments, while learned protein relationships could help recover information missing from individual panels and inform the choice of future measurements.

To address these linked challenges, we developed ProtCell, a foundation model for single-cell proteomics. We collected approximately 380 million cells from 98 datasets for pretraining and evaluation. ProtCell was pretrained using masked protein abundance reconstruction, enabling it to learn protein relationships shared across heterogeneous experimental panels and to generate representations at both the cell and protein levels. This shared representation supports analysis within individual datasets and connects different profiles through annotation transfer. Connecting cell states provides a basis for interpreting fragmented datasets, but the information available in each experiment remains restricted by its measured panel. We therefore draw on learned protein dependencies to extend profiles through abundance reconstruction and inference of proteins absent from target panels. These relationships also offer a way to guide future assay design by selecting compact panels that retain information about the broader protein profile. ProtCell thus links the analysis and connection of existing profiles with their extension and the design of more informative measurements.

## Results

We developed ProtCell, a protein language model for learning shared cell states and protein relationships across fragmented panels. We collected approximately 380 million cells from 98 datasets covering more than 600 protein targets for pretraining and downstream evaluation (Fig. 1a). Each cell forms a variable-length cell sentence, pairing protein identities with continuous abundances. A shared codebook supplies learned identity embeddings, while a multilayer perceptron encodes abundances without discretization. The codebook assigns the same identity to a protein across studies, allowing measurements from different panels to train its shared embedding. Combined identity and abundance embeddings enter a transformer alongside a cell-summary [CLS] token (Fig. 1b). Self-attention contextualizes each protein through the other measurements in that cell, accommodating different panels without restricting learning to their common protein intersection. We pretrained this language model with complementary protein- and cell-level reconstruction objectives. Masked abundance modeling retains protein identities while hiding selected measured values, requiring the model to predict abundances from their cellular context. Repeated masking across cells exposes each target to different combinations of available proteins, training predictions to draw on the measured context. A second head reconstructs the observed profile from [CLS], encouraging the cell summary to retain information distributed across proteins. Both objectives use measured targets and exclude unassayed values from the loss. Thus, heterogeneous panels contribute supervision for different parts of the shared vocabulary. The encoder produces contextual protein representations for profile completion and a cell representation for analysis and transfer, linking these tasks within one model (Fig. 1c).

**Figure 1.**
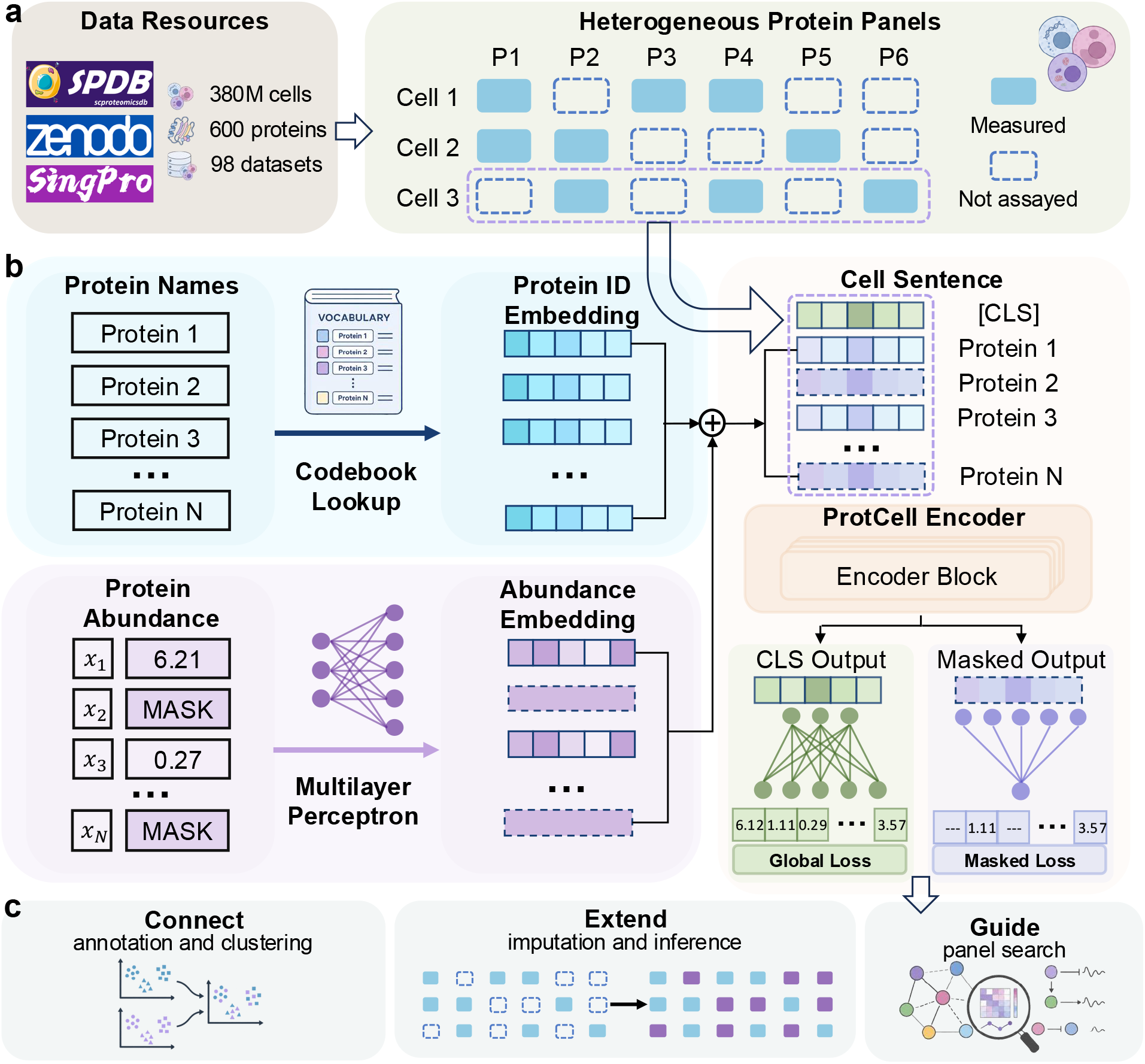
ProtCell learns shared representations across heterogeneous protein panels. **a**, Public single-cell proteomic resources were assembled into a collection of 98 datasets comprising approximately 380 million cells for pretraining and downstream evaluation. A schematic coverage matrix illustrates how heterogeneous panels measure partially overlapping protein sets, leaving different proteins unassayed across datasets. **b**, Protein identities are embedded through a shared codebook and continuous abundances through a multilayer perceptron. Their combined embeddings, together with a cell-level token, are processed by a transformer encoder. Cell-level and masked-position outputs are trained using global and masked reconstruction losses, respectively. **c**, The learned representations are evaluated for connecting profiles through annotation and clustering, extending profiles through protein imputation and inference, and guiding protein-panel search.

Limited assay coverage leaves informative proteins unmeasured, restricting the cell states and functional programs that each experiment can resolve. Profile extension therefore requires drawing on protein relationships learned beyond the target panel. We use masked language modeling with a shared codebook to learn these relationships across the atlas and condition predictions on each cell’s available measurements. This enables inference of proteins absent from the target panel but represented in the pretrained vocabulary. For these targets, the codebook supplies protein identities and the transformer supplies cellular context, reusing relationships learned in experiments where the targets were measured. Atlas pretraining makes these relationships reusable across target panels, whereas CytoVI’s imputation remains confined to its fitted marker set [13]. The identity-based vocabulary could also accommodate separately annotated modified forms, including phosphorylation-specific targets, with additional training.

We first benchmarked ProtCell through integration, annotation and masked protein imputation. In Beneyto-Calabuig2023, fine-tuning increased the weighted integration score from 0.33 zero-shot to 0.49, exceeding comparison methods [9] (Fig. 2a; Extended Data Fig. 1a–d). With the backbone frozen, within-batch annotation accuracy and macro F1 were approximately 0.98 in Kotliarov2020 [8] (Fig. 2b). Masked recovery improved over CytoVI in Peterson2021, Wagner2019 and Kotliarov2020 [7,8,13,17]; random-split mean protein-wise Pearson correlation in Kotliarov2020 was 0.586 versus 0.484 (Fig. 2d). Individually withholding proteins favoured ProtCell for 67 of 79 Kotliarov2020 targets by Pearson correlation (Fig. 2e; Extended Data Fig. 3a–c). Together, these benchmarks support both cell-level analysis and protein-level reconstruction.

**Figure 2.**
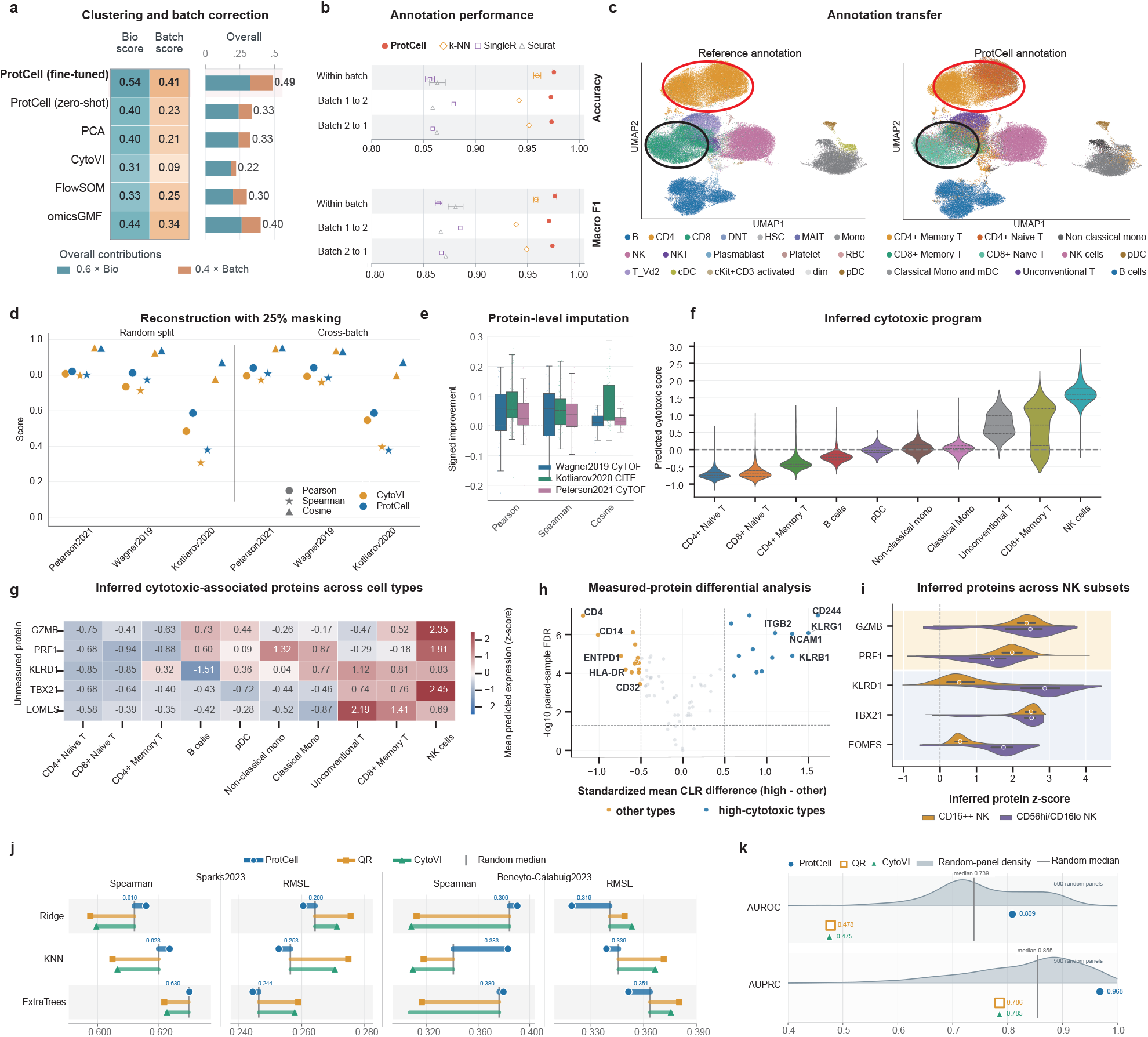
ProtCell connects and extends proteomic profiles and guides panel design. **a**, Average biological conservation, batch mixing and weighted overall integration scores for fine-tuned and zero-shot ProtCell and four comparison methods in Beneyto-Calabuig2023. Stacked bars show the contributions of 0.6 *×* avgBio and 0.4 *×* avgBatch to the overall score. **b**, Accuracy and macro F1 for cell-type annotation within batches and in two cross-batch transfer directions. **c**, UMAP representations of Sacco2022 coloured by its reference labels and by annotations transferred using the classifier trained on Kotliarov2020 with the ProtCell backbone frozen. **d**, Pearson correlation, Spearman correlation and cosine similarity after masking 25% of measured proteins in random-split and cross-batch settings. **e**, Protein-level distributions of the signed improvement of ProtCell over CytoVI from one-protein-at-a-time masking in Wagner2019, Kotliarov2020 and Peterson2021. **f**, Inferred cytotoxic score across immune cell types, calculated from GZMB, PRF1, KLRD1, TBX21 and EOMES. **g**, Standardized inferred abundances of the five unmeasured proteins across immune cell types. **h**, Paired-sample differential analysis of proteins measured in the target panel between cell types with high inferred cytotoxic scores and other cell types. **i**, Inferred distributions of the five unmeasured proteins in CD16-high and CD56-high/CD16-low natural killer-cell subsets. **j**, Mean protein-wise Spearman correlation and median protein-wise RMSE for panels selected by ProtCell-MSE, optimized anchor-constrained column-pivoted QR (QR) and CytoVI, evaluated using ridge, nearest-neighbour and extremely randomized tree decoders on common unmeasured targets in Sparks2023 and Beneyto-Calabuig2023, with medians from 50 random panels matched to the source budget. Source panels contain 60 markers; native mapping to Beneyto-Calabuig2023 retains 41, 39 and 40 markers, respectively, and 30–41 markers for random panels. **k**, Patient-median acute myeloid leukaemia discrimination using mapped 40-protein panels, compared with 500 size-matched random panels in three held-out patients.

We then tested transfer across batches and datasets. Kotliarov2020 cross-batch annotation accuracy remained 0.971–0.974, and mean protein-wise Pearson correlation for cross-batch imputation was 0.586 versus 0.546 for CytoVI (Fig. 2b,d). Transferring the annotation classifier to Sacco2022 distinguished naive and memory CD4^+^ and CD8^+^ T-cell states supported by measured markers [18–21] (Fig. 2c; Extended Data Fig. 2a–d). These results indicate that shared representations retain cell-state and protein information across experiments with different batches and panels.

We next tested whether ProtCell could extend a limited panel to reveal an unmeasured cytotoxic program. We inferred GZMB, PRF1, KLRD1, TBX21 and EOMES, absent from Kotliarov2020 but represented in the codebook [8, 22–26]. Their mean standardized inferred abundance defined a cytotoxic score, highest in natural killer cells (1.656) and lower in naive CD8^+^ T cells (*−*0.664) (Fig. 2f,g). Measured ITGB2, CD244, KLRG1 and NCAM1 were higher in high-scoring populations across ten paired samples; inferred natural killer-cell subset patterns agreed with reported biology [27–32] (Fig. 2h,i). This concordance supports using atlas-derived protein relationships to infer functional programs beyond the measured panel.

Finally, we explored whether learned protein relationships could guide measurements. With limited detection channels, panel search seeks protein combinations that maximize information about omitted targets. Frozen ProtCell reconstruction selected 60 of 128 proteins in Sparks2023. Across independent decoders, mean protein-wise Spearman correlations were 0.616–0.630 in Sparks2023 and 0.380–0.390 after transfer to Beneyto-Calabuig2023, with lower RMSE than comparison panels [9, 33] (Fig. 2j). Mapped 40-protein panels yielded median patient-level AUROC/AUPRC of 0.809/0.968 for acute myeloid leukaemia clone discrimination in three held-out patients, versus 0.478/0.786 for QR and 0.475/0.785 for CytoVI (Fig. 2k). This exploratory application connects profile extension to informative assay design.

## Discussion

ProtCell brings information distributed across fragmented proteomic panels into shared cell- and protein-level representations. The codebook establishes consistent protein identities across studies, while masked language modeling learns relationships conditional on cellular context. Together, they support annotation transfer, reconstruction and inference beyond individual panels, linking the interpretation of existing measurements to the design of future assays. The inferred cytotoxic program illustrates how profile extension can add a functional perspective to cell identity, with predictions supported by measured markers and known lymphocyte biology. Panel search further suggests that these relationships can help retain useful information within limited detection channels.

Current protein coverage remains concentrated on immune-associated targets, and modified forms are not yet systematically distinguished, reflecting gaps in available training data. Broader datasets with explicit modification annotations could extend the codebook beyond immune panels and distinguish targets such as phosphorylated forms. Incorporating protein interaction and pathway knowledge could further inform representations when measured context is sparse. Panel search could also be developed for specific biological objectives and practical assay constraints. Together with direct measurements of inferred targets and prospective panel validation, these directions would broaden ProtCell’s capacity to connect fragmented profiles, extend their functional coverage and guide informative measurements.

## Methods

### Data collection and protein vocabulary construction

Public single-cell proteomic datasets were collected from SPDB, Zenodo-hosted resources, SingPro, and related public repositories [34, 35]. Datasets were manually reviewed to remove entries with incompatible formats, insufficient metadata, poor quality, or ambiguous protein annotations. The collection comprises approximately 380 million cells from 98 datasets. The retained pretraining inputs and their cell and target counts are detailed in Supplementary Table 1.

Protein names were standardized into a shared protein vocabulary. In the model implementation, `ProteinTokenizer` maps protein names to integer token identifiers and maintains special tokens including `<pad>` and `<eos>`. This protein vocabulary allows ProtCell to process datasets with different measured panels by representing proteins through shared token identities rather than fixed matrix columns.

### Preprocessing and abundance standardization

Protein abundance matrices were transformed before modeling to reduce platform-specific scale differences. CITE-seq measurements were represented with centered log-ratio transformations (CLR), whereas CyTOF measurements used arcsinh transformations (labelled ARCSIN in the metadata). The training reader retained only entries matching these technology-specific preprocessing labels. Missing or unmeasured proteins were not treated as true zero abundance. In dataloaders, invalid or padded positions were represented with sentinel values and excluded from observed-position losses. During masked modeling, selected observed abundances were replaced by a learned mask embedding.

### ProtCell architecture

ProtCell encodes each cell as a variable-length set of protein tokens paired with standardized abundance values. Protein identities are mapped to discrete token identifiers through a protein codebook and embedded in a shared latent space, whereas continuous abundance values are encoded by a multilayer perceptron. The identity and abundance embeddings are combined as token-level inputs to a transformer encoder. A learnable cell-level token summarizes the full proteomic profile for downstream analyses, while token-level outputs are decoded through a regression head to predict protein abundance values.

### Pretraining and masked abundance objective

The core training objective is masked abundance reconstruction [36]. For each cell, a fraction of observed protein positions is randomly selected for masking. The model receives the protein identities and the partially masked abundance profile, and predicts the held-out abundance values. The main loss is mean squared error over masked observed positions:

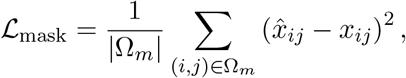

where Ω_*m*_ denotes masked observed positions. An additional reconstruction term is computed over observed profile values selected by input protein identifiers. In the current implementation this term is added to the masked reconstruction loss when available.

We also reproduced ProtCell on Moore Threads MTT S5000 GPUs, obtaining broadly comparable performance with approximately 0.2% variation in quantitative evaluation metrics.

### Cell embedding extraction

For downstream cell-level tasks, ProtCell uses the projected CLS representation as the default cell embedding. The implementation also supports mean pooling, but the Results described here are framed around CLS embeddings. These embeddings are used for cell-type annotation, clustering, integration, and batch-correction fine-tuning.

### Cell-type annotation benchmark

Cell-type annotation was evaluated in intra-batch, cross-batch and cross-dataset settings. The entire ProtCell backbone was frozen and used only to extract CLS embeddings; a multilayer classification head was trained with cross-entropy loss. In intra-batch experiments, cells were split into five folds within the same batch. In cross-batch experiments, the classification head was trained on one batch and evaluated on another. For cross-dataset transfer, the head trained using the more resolved Kotliarov2020 annotations was evaluated in Sacco2022. An optional class-weighted version uses inverse square-root class-frequency weights to reduce the influence of highly abundant cell types.

ProtCell was compared with *k*-nearest neighbours, SingleR [37], and Seurat [38, 39]. Metrics were computed from classification reports and summarized as accuracy, macro F1, and weighted F1. For intra-batch experiments, fold-level values were summarized as mean performance with variability. For cross-batch experiments, each train-to-test direction was treated as a transfer experiment rather than a replicate distribution.

### Batch correction and integration fine-tuning

Integration fine-tuning uses ProtCell CLS embeddings and an unbalanced optimal transport-based contrastive objective. The adapter samples a target batch within a mini-batch and treats cells from the remaining batches as potential cross-batch counterparts. A cosine-distance cost matrix is computed between target and rest embeddings. Related batch-integration frameworks have shown the value of explicitly modeling cross-batch structure in single-cell data [40, 41]. An unbalanced Sinkhorn optimal transport plan is estimated, allowing unequal cell-state composition across batches [42, 43]. The resulting transport weights define cross-batch positive relationships in a contrastive loss.

To preserve local structure within batches, an additional intra-batch distribution preservation term compares the current within-batch similarity distribution with a reference embedding distribution. The total integration loss combines the transport-weighted contrastive term and the intra-batch preservation term. LoRA adapters can be deployed on selected transformer and CLS projection modules for parameter-efficient fine-tuning [44], and merged back into the model when saving.

### Clustering and integration evaluation

Clustering and integration were evaluated on Sacco2022 and Beneyto-Calabuig2023 [9, 18]. Methods compared included zero-shot ProtCell, fine-tuned ProtCell, PCA, CytoVI, FlowSOM and omicsGMF [12–14]. Metrics included the adjusted Rand index, normalized mutual information, silhouette-based scores, kBET, iLISI, Bio_ASW, Batch_ASW, avgBio and avgBatch [45]. The figure-level overall score used

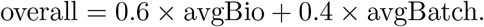

Detailed integration and annotation results are described in the Supplementary Note 1 on connecting fragmented profiles.

### Masked protein imputation benchmark

Protein abundance imputation was evaluated against CytoVI [13] using Wagner2019, Kotliarov2020 and Peterson2021 [7,8,17]. ProtCell was fine-tuned within each dataset to adapt to its measurement distribution. In the random half-cell design, one half of the cells was used for fine-tuning and the other half for testing. In the cross-batch or cross-condition design, fine-tuning used one batch or condition and evaluation used the remaining batches or conditions. In the 25% masking setting, a quarter of measured proteins was randomly masked and reconstructed. In the protein-wise setting, each protein was masked and predicted one at a time to assess marker-level recovery. Metrics included Pearson correlation, Spearman correlation, cosine similarity, normalized mean squared error, mean squared error, normalized mean absolute error and mean absolute error.

For paired comparison, signed improvement was oriented so that positive values always indicate ProtCell advantage. For higher-is-better metrics,

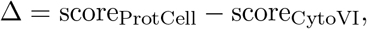

whereas for lower-is-better error metrics,

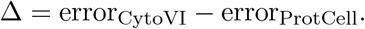

Win rate was defined as the fraction of paired comparisons for which Δ *>* 0. Detailed reconstruction results are described in the Supplementary Note 2 on recovering measured proteins.

### Inference of unmeasured T/NK effector markers

For Kotliarov2020 [8], candidate unmeasured proteins were selected from the ProtCell vocabulary among proteins absent from the measured panel but biologically relevant to immune effector states. The final marker set comprised GZMB, PRF1, KLRD1, TBX21 and EOMES. Predicted values were analysed together with the measured protein abundances and original cell metadata.

A cytotoxic score was computed per cell by z-scoring each predicted marker across all cells and averaging the standardized values:

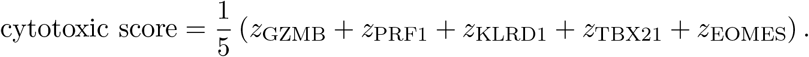

Measured-protein support for the inferred program was evaluated at the biological-sample level. Centered log-ratio abundances of measured proteins were summarized separately for cell types with high inferred cytotoxic scores and for the remaining cell types, and standardized paired-sample differences were calculated. Natural killer-cell subtype analysis included 2,460 cells from ten paired biological samples. Cell-level distributions and paired sample-level differences in inferred protein abundance were compared between CD16-high and CD56-high/CD16-low natural killer-cell subsets. Detailed biological support is described in the Supplementary Note 3 on the unmeasured cytotoxic program.

### Protein-panel selection: objective and search procedure

Protein-panel selection seeks a fixed-size set of measurements that retains information about the broader protein profile while including markers for major cell lineages. We use “panel search” to describe optimization of this set for reconstruction of unselected proteins. Existing complete measurements allowed retrospective masking and scoring of candidate panels; this tests information retention rather than the experimental performance of a new antibody panel. Sparks2023 [33] provided 128 candidates, with 16 discovery, eight selection and 16 test donors. All source panels contained 60 markers, including six mandatory anchors: CD3, CD4, CD19, CD14, KLRD1 and the combined FCGR3A/FCGR3B measurement.

For ProtCell-based selection, the pretrained checkpoint was fixed throughout search. Given a panel *S*, all remaining proteins *T* were masked simultaneously and predicted from the selected measurements. Candidate error was

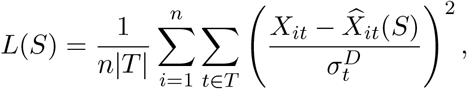

where 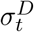 was the protein standard deviation in discovery cells; scales at or below 10^*−*8^ were replaced by one. Twenty cell resamples were crossed with ten search seeds, yielding 200 runs. Each resample drew 32 cells without replacement from each selection donor–visit stratum (768 cells), leaving 6,432 selection cells for evaluating run winners. Each run scored 40 random anchored panels, then refined the best through two rounds of 24 single-marker exchange proposals, retaining the lowest-error panel at each round. Exchanges preserved all anchors and the marker budget. The final panel minimized winner error on the remaining selection cells, which came from the same selection donors and were separate from the test cohort.

### Anchor-constrained column-pivoted QR selection

The QR baseline used column-pivoted QR (CPQR), a classical matrix column-selection approach [46]. Unlike ordinary QR, CPQR reorders columns, greedily favouring proteins with large residual variation after projection onto previously selected columns. We first centered and standardized each protein column of the discovery expression matrix *Z* using population standard deviations, replacing scales at or below 10^*−*8^ by one. For anchor columns *A* and optional columns *O*, we computed a reduced QR factorization *Z*_*A*_ = *Q*_*A*_*R*_*A*_, followed by

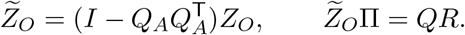

Here *Q*_*A*_ spans the anchor directions and Π encodes the optional-column pivot order. The first 54 pivots were combined with the six anchors. This prioritizes additional linear information beyond the mandatory markers. The approach is related to QR-based sparse sensor placement [47], but operates directly on standardized protein-expression columns without first constructing a truncated SVD/POD basis.

The optimized QR comparator shown in the main figure repeated initialization over 50 resamples, each drawing 250 discovery cells without replacement from each donor–visit stratum (12,000 cells per run). Each resulting panel underwent up to three single-marker swaps, with anchors protected. Refinement used ridge-linear reconstruction of omitted proteins: coefficients were fitted from discovery data with a penalty of 10^*−*4^ added to the diagonal of the sample-normalized predictor second-moment matrix. Panels were scored by mean standardized reconstruction MSE on selection donors, using fixed discovery means and scales. First-improvement and exhaustive best-improvement searches were evaluated, including an exhaustive variant that recorded all candidate scores; the lowest-scoring result across strategies and runs was retained. Thus, randomness arose from cell resampling, and the displayed QR comparator included reconstruction-guided refinement beyond CPQR initialization.

### CytoVI selection and freezing of panels

CytoVI-based search [13] used 20 starting panels comprising two earlier reference panels, nine diverse optimized QR panels and nine anchored random panels. Each start underwent two rounds of 24 exchange proposals, with eight each from uniform, reconstruction-error and expression-structure-guided proposals. Each candidate CytoVI model was fitted on discovery donors represented as complete-panel and candidate-panel batches, with discovery-derived standardization. Models were trained for up to 100 epochs with a batch size of 512 and an early-stopping patience of 15 epochs. Candidates were ranked by standardized reconstruction RMSE on selection donors. Each run winner was retrained with five seeds, and the panel with the lowest median selection RMSE was retained.

The main figure compares one fixed panel per selector, chosen using its own selection objective; objective values were not compared across selectors. ProtCell, QR and CytoVI used 200, 50 and 20 search runs, respectively, so this comparison does not match search budgets. Separate matched-best-of-20 analyses evaluated run-count matching. Test-cohort reconstruction results and AML labels were not used to score or select panels. Historical test results had nevertheless already been inspected during earlier analyses, so the evaluation is retrospective rather than a newly blinded confirmatory test.

### Independent panel-completion evaluation

To assess retained information using predictors distinct from the selection models, we fitted ridge regression, *k*-nearest-neighbour regression and extremely randomized trees to each panel. Evaluation used identical test cells and proteins absent from all three selected panels. For each dataset, 50 random reference panels retained the anchors, matched the source budget of 60 markers and excluded the corresponding common targets. Sparks2023 evaluation used 39 common targets and 24,000 test cells. For transfer to Beneyto-Calabuig2023 [9], native mapping retained 41, 39 and 40 measured proteins from the ProtCell, QR and CytoVI panels, respectively. Marker sets were transferred, while decoders were fitted using Cohort A and tested on 45,903 Cohort B cells over 80 common targets. Random panels retained 30–41 markers after mapping. Thus, this completion comparison matched source budgets but retained unequal mapped panel sizes.

For each target protein, Spearman correlation and RMSE were computed across test cells; the displayed summaries were the mean correlation and median RMSE across targets. RMSE used the expression values supplied to the decoders, without division by discovery protein scales, and therefore differs from the standardized search objective. The three decoders and two protein-wise metrics in the main figure were chosen after inspection of available results; broader decoder and global-metric results were retained separately.

### Acute myeloid leukaemia panel evaluation

To test whether the selected measurements also retained information relevant to a biological classification task, we evaluated acute myeloid leukaemia clone discrimination. Each selected panel was mapped to exactly 40 proteins measured in Beneyto-Calabuig2023 [9]. Panel mapping and marker backfilling followed the original candidate-protein order without using leukaemia labels: excess mapped markers were truncated, and missing markers were filled from available source candidates. This fixed-size comparison was separate from the completion analysis, which retained the different panel sizes after native mapping. Only these 40 measured proteins were used as classifier inputs; reconstructed protein values were not included. A logistic regression classifier was trained using Cohort A and evaluated separately in three held-out Cohort B patients, with up to 1,000 labelled myeloid cells sampled per patient and class. Performance was calculated within each patient and summarized using the median area under the receiver operating characteristic curve and median area under the precision–recall curve. The QR and CytoVI panels were evaluated using the same procedure, and 500 matched random 40-protein panels provided a computational reference. Detailed completion and classification results are described in the Supplementary Note 4 on protein-panel search.

## Code availability

The ProtCell source code for pretraining and task-specific adaptation, together with downstream analysis notebooks, is available at https://github.com/hubwenbwen/ProtCell. Installation instructions and example workflows are provided in the repository.

## Extended Data Figures

**Extended Data Fig. 1.**
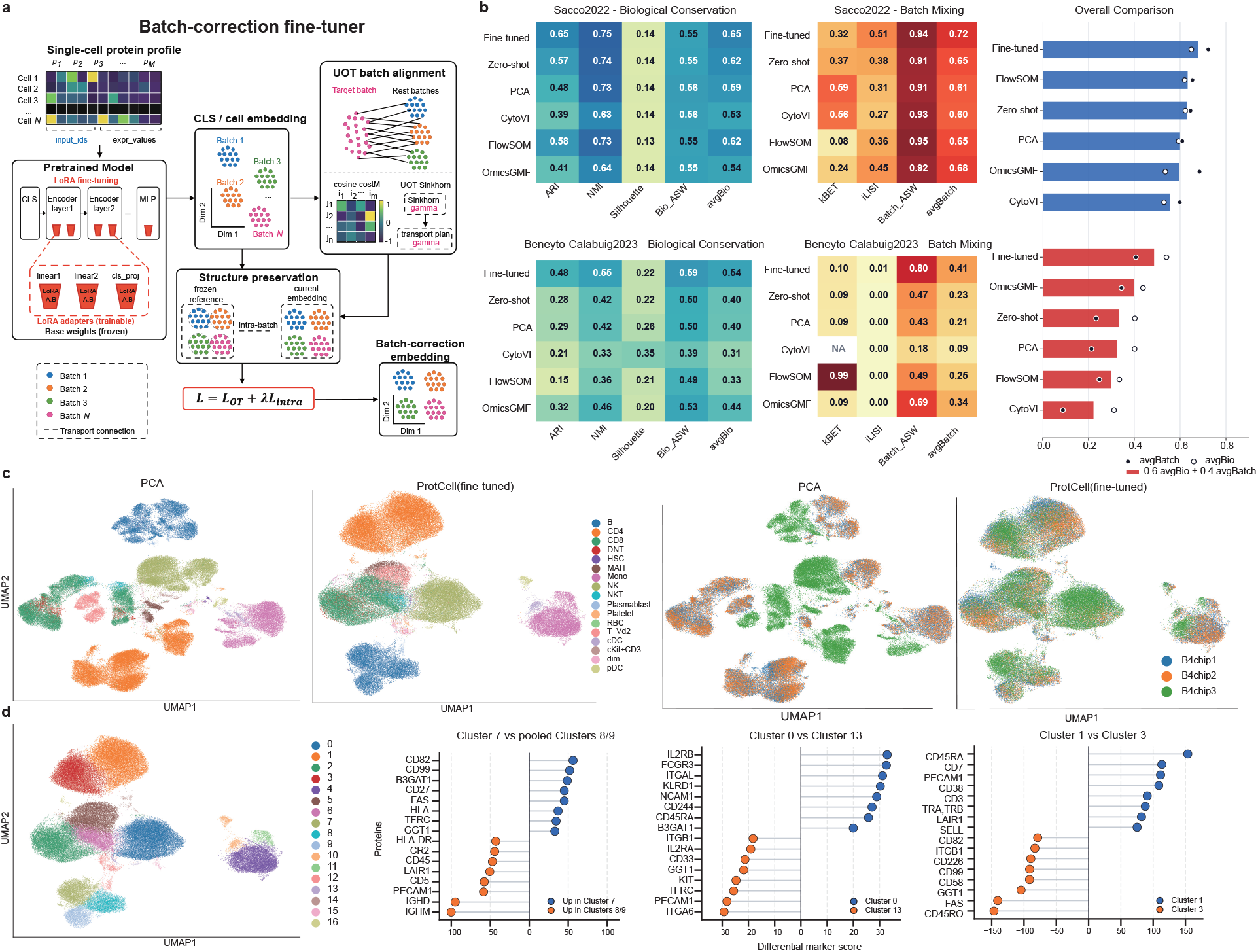
Batch-correction design, complete integration benchmarks and cluster-level protein states. **a**, Contrastive fine-tuning aligns cell embeddings across batches using unbalanced optimal transport, while preserving within-batch structure; the pretrained back-bone remains frozen and LoRA adapters are trained. **b**, Biological conservation and batch-mixing metrics in Sacco2022 and Beneyto-Calabuig2023 for fine-tuned and zero-shot ProtCell and the comparison methods, together with weighted overall scores. **c**, Sacco2022 UMAPs from PCA and fine-tuned ProtCell representations, coloured by reference cell type (left) and experimental batch (right). **d**, Cluster assignments in the adapted Sacco2022 representation and differential measured-protein patterns across three displayed cluster contrasts. Positive and negative scores indicate enrichment in the respective clusters named in each contrast.

**Extended Data Fig. 2.**
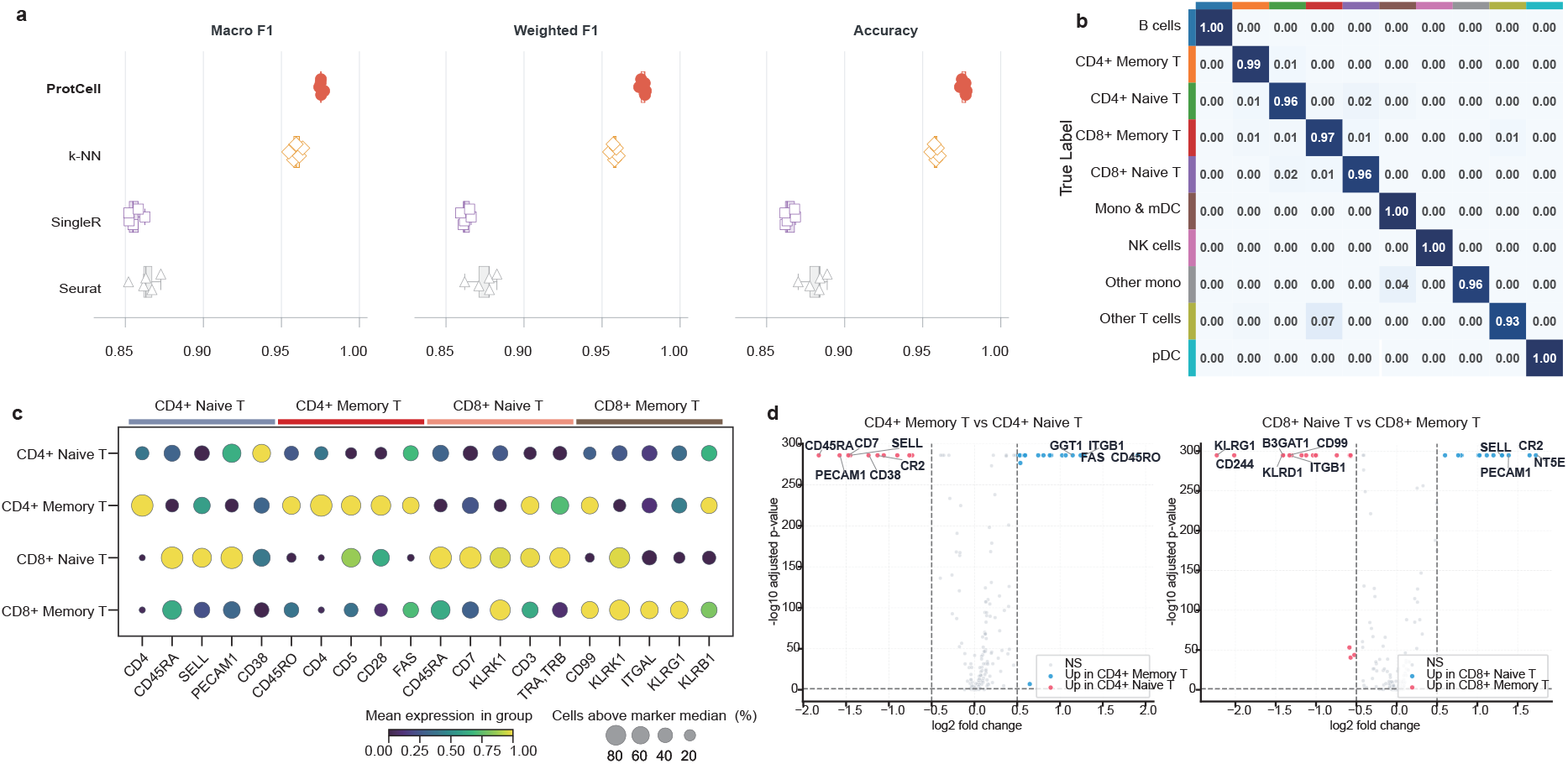
Detailed annotation transfer and measured-protein support. **a**, Macro F1, weighted F1 and accuracy for ProtCell, *k*-nearest neighbours, SingleR and Seurat in the Kotliarov2020 annotation benchmark across within-batch and bidirectional cross-batch settings. **b**, Cross-batch confusion matrix for major immune populations, with rows representing reference labels. **c**, Measured-protein patterns across transferred naive and memory CD4^+^ and CD8^+^ T-cell annotations in Sacco2022. Dot size shows the percentage of cells above the marker median, and colour shows mean expression within each group. **d**, Differential measured-protein signals for naive and memory T-cell contrasts within CD4^+^ and CD8^+^ populations.

**Extended Data Fig. 3.**
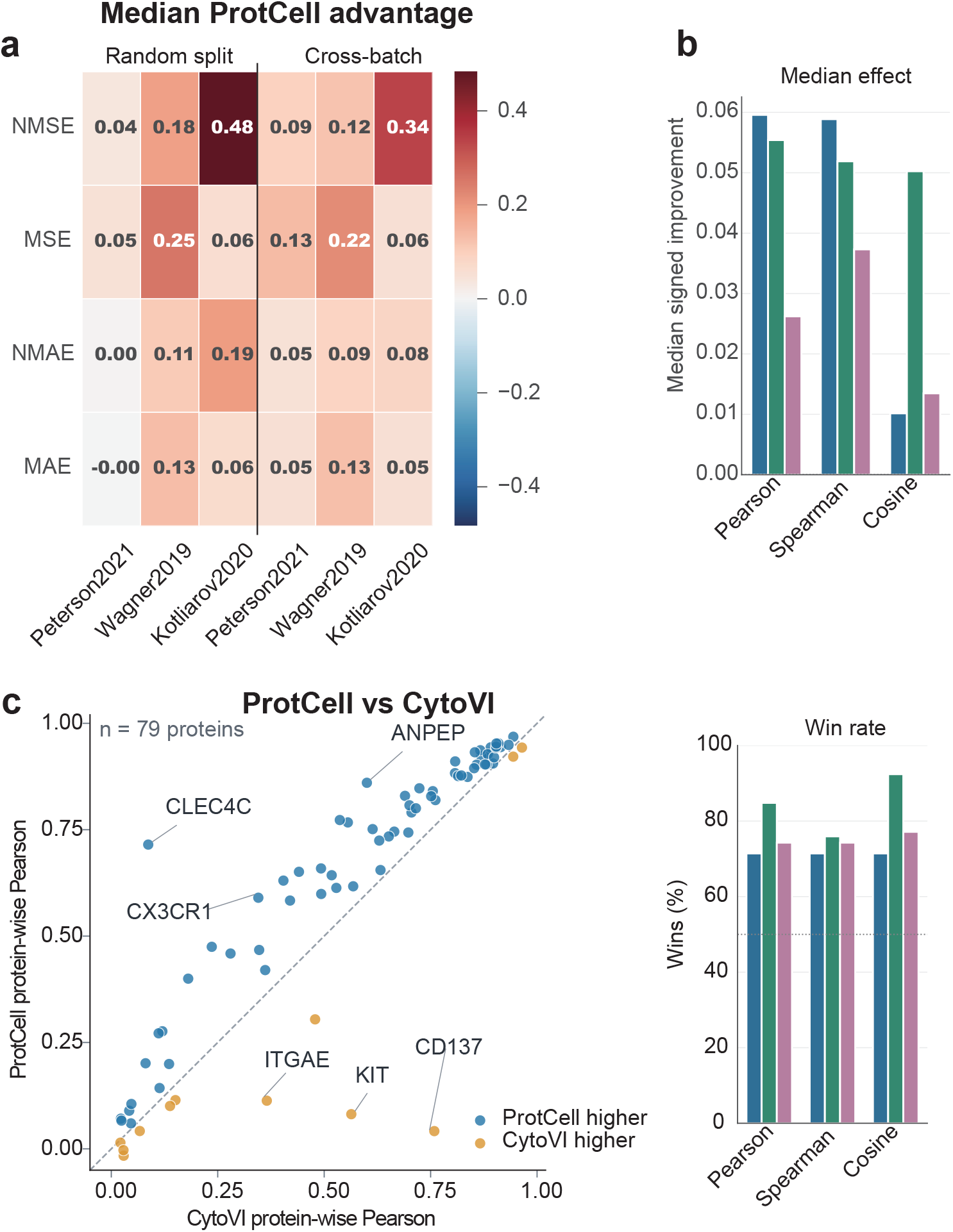
Additional imputation comparisons with CytoVI. **a**, Median Prot-Cell advantage over CytoVI for normalized mean squared error, mean squared error, normalized mean absolute error and mean absolute error across three datasets in random-split and cross-batch settings. **b**, Median signed protein-wise improvements by Pearson correlation, Spearman correlation and cosine similarity after one-protein-at-a-time masking. **c**, Protein-wise Pearson correlations for ProtCell and CytoVI in Kotliarov2020, with the diagonal indicating equal performance, and protein-wise win rates for all three datasets and three similarity measures.

## References

[1] Ruedi Aebersold and Matthias Mann. Mass-spectrometric exploration of proteome structure and function. Nature, 537:347–355, 2016.

[2] Björn Schwanhäusser et al. Global quantification of mammalian gene expression control. Nature, 473:337–342, 2011.

[3] Yansheng Liu, Andreas Beyer, and Ruedi Aebersold. On the dependency of cellular protein levels on mRNA abundance. Cell, 165:535–550, 2016.

[4] Hayley M. Bennett, William Stephenson, Christopher M. Rose, and Spyros Darmanis. Single-cell proteomics enabled by next-generation sequencing or mass spectrometry. Nature Methods, 20:363–374, 2023.

[5] Sophia M. Guldberg, Trine Line Hauge Okholm, Elizabeth E. McCarthy, and Matthew H. Spitzer. Computational methods for single-cell proteomics. Annual Review of Biomedical Data Science, 6:47–71, 2023.

[6] Sean C. Bendall et al. Single-cell mass cytometry of differential immune and drug responses across a human hematopoietic continuum. Science, 332:687–696, 2011.

[7] Johanna Wagner et al. A single-cell atlas of the tumor and immune ecosystem of human breast cancer. Cell, 177:1330–1345.e18, 2019.

[8] Yuri Kotliarov et al. Broad immune activation underlies shared set point signatures for vaccine responsiveness in healthy individuals and disease activity in patients with lupus. Nature Medicine, 26:618–629, 2020.

[9] Sergi Beneyto-Calabuig et al. Clonally resolved single-cell multi-omics identifies routes of cellular differentiation in acute myeloid leukemia. Cell Stem Cell, 30:706–721.e8, 2023.

[10] Dmitry R. Bandura et al. Mass cytometry: technique for real time single cell multitarget immunoassay based on inductively coupled plasma time-of-flight mass spectrometry. Analytical Chemistry, 81:6813–6822, 2009.

[11] Marlon Stoeckius et al. Simultaneous epitope and transcriptome measurement in single cells. Nature Methods, 14:865–868, 2017.

[12] Sofie Van Gassen et al. Flowsom: using self-organizing maps for visualization and interpretation of cytometry data. Cytometry Part A, 87:636–645, 2015.

[13] Florian Ingelfinger, Nathan Levy, Can Ergen, et al. CytoVI: deep generative modeling of antibody-based single cell data. Nature Methods, 23:2054–2065, 2026.

[14] Alexandre Segers et al. omicsgmf: a multi-tool for dimensionality reduction, batch correction and imputation in bulk- and single-cell proteomics. Nature Communications, 17:6650, 2026.

[15] Haotian Cui et al. scgpt: toward building a foundation model for single-cell multi-omics using generative ai. Nature Methods, 21:1470–1480, 2024.

[16] Minsheng Hao et al. Large-scale foundation model on single-cell transcriptomics. Nature Methods, 21:1481–1491, 2024.

[17] Laura S. Peterson et al. Single-cell analysis of the neonatal immune system across the gestational age continuum. Frontiers in Immunology, 12:714090, 2021.

[18] Keith Sacco et al. Immunopathological signatures in multisystem inflammatory syndrome in children and pediatric covid-19. Nature Medicine, 28:1050–1062, 2022.

[19] Federica Sallusto, Danielle Lenig, Reinhold Förster, Martin Lipp, and Antonio Lanzavecchia. Two subsets of memory t lymphocytes with distinct homing potentials and effector functions. Nature, 401:708–712, 1999.

[20] Luca Gattinoni et al. A human memory t cell subset with stem cell-like properties. Nature Medicine, 17:1290–1297, 2011.

[21] N. K. Damle and L. V. Doyle. Ability of human T lymphocytes to adhere to vascular endothelial cells and to augment endothelial permeability to macromolecules is linked to their state of post-thymic maturation. The Journal of Immunology, 144(4):1233–1240, 1990.

[22] John H. Russell and Timothy J. Ley. Lymphocyte-mediated cytotoxicity. Annual Review of Immunology, 20:323–370, 2002.

[23] Ilia Voskoboinik, James C. Whisstock, and Joseph A. Trapani. Perforin and granzymes: function, dysfunction and human pathology. Nature Reviews Immunology, 15:388–400, 2015.

[24] Eric Vivier, Elena Tomasello, Myriam Baratin, Thierry Walzer, and Sophie Ugolini. Functions of natural killer cells. Nature Immunology, 9:503–510, 2008.

[25] Susan M. Kaech and Weiguo Cui. Transcriptional control of effector and memory cd8+ t cell differentiation. Nature Reviews Immunology, 12:749–761, 2012.

[26] Scott M. Gordon et al. The transcription factors t-bet and eomes control key checkpoints of natural killer cell maturation. Immunity, 36:55–67, 2012.

[27] Jordan S. Orange. Formation and function of the lytic NK-cell immunological synapse. Nature Reviews Immunology, 8(9):713–725, 2008.

[28] Nikhil S. Joshi et al. Inflammation directs memory precursor and short-lived effector CD8+ T cell fates via the graded expression of T-bet transcription factor. Immunity, 27(2):281–295, 2007.

[29] Michael A. Caligiuri. Human natural killer cells. Blood, 112(3):461–469, 2008.

[30] Jianhua Yu et al. CD94 surface density identifies a functional intermediary between the CD56bright and CD56dim human NK-cell subsets. Blood, 115(2):274–281, 2010.

[31] James J. Knox, Gabriela L. Cosma, Michael R. Betts, and Laura M. McLane. Characterization of T-bet and Eomes in peripheral human immune cells. Frontiers in Immunology, 5:217, 2014.

[32] Julia A. Wagner et al. CD56bright NK cells exhibit potent antitumor responses following IL-15 priming. The Journal of Clinical Investigation, 127(11):4042–4058, 2017.

[33] Rachel Sparks et al. Influenza vaccination reveals sex dimorphic imprints of prior mild COVID-19. Nature, 614(7949):752–761, 2023.

[34] Fang Wang et al. Spdb: a comprehensive resource and knowledgebase for proteomic data at the single-cell resolution. Nucleic Acids Research, 52:D562–D571, 2024.

[35] Xichen Lian et al. Singpro: a knowledge base providing single-cell proteomic data. Nucleic Acids Research, 52:D552–D561, 2024.

[36] Jacob Devlin, Ming-Wei Chang, Kenton Lee, and Kristina Toutanova. Bert: pre-training of deep bidirectional transformers for language understanding. In Proceedings of NAACL-HLT, pages 4171–4186, 2019.

[37] Dvir Aran et al. Reference-based analysis of lung single-cell sequencing reveals a transitional profibrotic macrophage. Nature Immunology, 20:163–172, 2019.

[38] Tim Stuart et al. Comprehensive integration of single-cell data. Cell, 177:1888–1902.e21, 2019.

[39] Yuhan Hao et al. Integrated analysis of multimodal single-cell data. Cell, 184:3573–3587.e29, 2021.

[40] Ilya Korsunsky et al. Fast, sensitive and accurate integration of single-cell data with harmony. Nature Methods, 16:1289–1296, 2019.

[41] Christina Bligaard Pedersen et al. cycombine allows for robust integration of single-cell cytometry datasets within and across technologies. Nature Communications, 13:1698, 2022.

[42] Marco Cuturi. Sinkhorn distances: lightspeed computation of optimal transport. In Advances in Neural Information Processing Systems, volume 26, 2013.

[43] Lénaïc Chizat, Gabriel Peyré, Bernhard Schmitzer, and François-Xavier Vialard. Scaling algorithms for unbalanced optimal transport problems. Mathematics of Computation, 87:2563–2609, 2018.

[44] Edward J. Hu et al. Lora: low-rank adaptation of large language models. In International Conference on Learning Representations, 2022.

[45] Malte D. Luecken et al. Benchmarking atlas-level data integration in single-cell genomics. Nature Methods, 19:41–50, 2022.

[46] Peter Businger and Gene H. Golub. Linear least squares solutions by Householder transformations. Numerische Mathematik, 7:269–276, 1965.

[47] Krithika Manohar, Bingni W. Brunton, J. Nathan Kutz, and Steven L. Brunton. Data-driven sparse sensor placement for reconstruction: Demonstrating the benefits of exploiting known patterns. IEEE Control Systems Magazine, 38(3):63–86, 2018.

