## Supplementary Information for "ProtCell: a foundation model for single-cell proteomics"

This document contains Supplementary Notes 1–4 and Supplementary Table 1.

### Supplementary Notes

#### Supplementary Note 1. Connecting fragmented profiles through cell representations

We first asked whether ProtCell could provide a shared description of cell states across fragmented proteomic experiments. Differences in measured proteins and experimental batches can obscure biological relationships between cells, even when studies sample related populations. We therefore examined two complementary uses of the cell representation: integration and clustering after adaptation, and annotation with the pretrained backbone frozen. The former tests whether cell structure can be aligned across batches; the latter tests whether the pretrained representation already contains information that transfers to an annotation task. Together, these analyses examine whether shared representations can link cell states across fragmented proteomic experiments.

For integration, we adapted the cell representation using transport-based cross-batch alignment while preserving within-batch structure (Extended Data Fig. 1a). In Beneyto-Calabuig2023 [9], the weighted overall score increased from 0.335 without adaptation to 0.487 after fine-tuning. Biological conservation increased from 0.402 to 0.540, and batch mixing from 0.233 to 0.407. The overall scores for PCA, CytoVI, FlowSOM and omicsGMF were 0.326, 0.222, 0.300 and 0.400, respectively [12–14] (Fig. 2a). Fine-tuning improved both biological conservation and batch mixing. In Sacco2022 [17], the same weighted score increased from 0.632 to 0.679, with biological conservation increasing from 0.623 to 0.649 and batch mixing from 0.645 to 0.723 (Extended Data Fig. 1b). The UMAP and cluster-level marker analyses provide complementary views of the retained cell organization (Extended Data Fig. 1c,d).

For annotation, we trained a classification head on frozen ProtCell embeddings in Kotliarov2020 [8]. Across five within-batch folds, mean accuracy was 0.976 and macro F1 was 0.976, with sample standard deviations of 0.002 and 0.001. Accuracy and macro F1 were 0.958 and 0.959 for $k$-nearest neighbours, 0.865 and 0.856 for SingleR, and 0.881 and 0.863 for Seurat [37,38] (Fig. 2b; Extended

Data Fig. 2a). Macro F1 complements accuracy by summarizing classification performance across cell types. These results support the use of the pretrained cell representation for annotation through a classification head trained on frozen embeddings.

We then transferred annotation between the two Kotliarov2020 batches. Accuracy was 0.971 for batch 1 to batch 2 and 0.974 in the reverse direction, compared with 0.939 and 0.949 for $k$-nearest neighbours. ProtCell macro F1 remained approximately 0.973 in both directions, compared with 0.942 and 0.952 for $k$-nearest neighbours. Thus, the annotation advantage persisted under the batch shift and involved both overall classification and performance across cell types.

Cross-dataset transfer to Sacco2022 tested a further step towards connecting experiments with differing panel compositions. The Kotliarov2020-trained classifier retained broad cell organization while distinguishing naive and memory $CD4^+$ and $CD8^+$ T-cell states (Fig. 2c). Measured CD45RA, SELL, CD45RO and other lineage-associated markers supported these distinctions [18–20] (Extended Data Fig. 2c,d). Together, the integration and transfer results show how a shared cell representation can connect fragmented measurements. Having connected cell states across experiments, we next asked whether protein representations could extend the coverage of individual panels.

#### Supplementary Note 2. Extending incomplete profiles by recovering measured proteins

We next tested whether learned protein context could recover information removed from measured profiles. Masking observed proteins provides a directly testable imputation task because the original abundances remain available as ground truth. We evaluated Peterson2021, Wagner2019 and Kotliarov2020 [7, 8, 21], comparing ProtCell with CytoVI after dataset-specific training [13]. Half-cell splits tested recovery within a dataset, whereas transfer across batches or conditions tested recovery under a change in measurement distribution. In the approximately 25% masking setting, protein-wise correlations were calculated across test cells and averaged over the masked targets. These summaries capture the recovery of individual protein patterns across cells.

For the half-cell split, mean protein-wise Pearson correlation was 0.821 for ProtCell versus 0.808 for CytoVI in Peterson2021, 0.812 versus 0.735 in Wagner2019, and 0.586 versus 0.484 in Kotliarov2020 (Fig. 2d). The magnitude of improvement varied across datasets: the difference was modest in Peterson2021 and larger in Wagner2019 and Kotliarov2020. In Wagner2019, mean Spearman correlation also increased from 0.713 to 0.773. In Kotliarov2020, it increased from 0.307 to 0.379, while mean normalized mean squared error decreased from 1.069 to 0.586 and normalized mean absolute error from 0.726 to 0.534. These comparisons support recovery of both cross-cell abundance patterns and numerical values.

We also tested cross-batch or cross-condition imputation. For example, mean protein-wise Pearson correlation reached 0.841 versus 0.796 in Peterson2021 and 0.586 versus 0.546 in Kotliarov2020. In Kotliarov2020, cross-batch cosine similarity was 0.873 versus 0.798, and Spearman correlation was 0.377 versus 0.396. Together with the error measures, these metrics characterize complementary aspects of abundance recovery (Fig. 2d; Extended Data Fig. 3a).

We also tested recovery by withholding each protein individually. In the Kotliarov2020 cross-batch comparison, ProtCell exceeded CytoVI for 67 of 79 targets by Pearson correlation (84.8%), 60 by Spearman correlation and 73 by cosine similarity. Protein-wise signed differences show the direction and distribution of improvement, while the paired Pearson plot identifies proteins for which either method performed better (Fig. 2e; Extended Data Fig. 3b,c). These results support the recovery of missing measurements across many targets through contextual protein representations. We then extended the analysis to proteins absent from the target panel.

#### Supplementary Note 3. Extending fragmented panels to an unmeasured cytotoxic program

We then asked whether ProtCell could infer a biological program whose constituent proteins were absent from the target panel. We selected GZMB, PRF1, KLRD1, TBX21 and EOMES, which were represented in the pretrained vocabulary but unmeasured in Kotliarov2020 [8]. GZMB and PRF1 are cytolytic effectors, KLRD1 encodes the CD94 receptor, and TBX21 and EOMES are transcription factors associated with effector lymphocyte differentiation [22–26]. After fine-tuning on batch 1, we examined inference in batch 2, containing 24,972 cells from ten biological samples and 79 measured proteins. This setting tests extension of the target assay's coverage through protein relationships learned elsewhere in the atlas.

The mean of the five standardized inferred abundances defined a cytotoxic score. The reported cell-type means were 1.656 in natural killer cells, 0.720 in unconventional T cells and 0.652 in memory $CD8^+$ T cells, compared with $-0.664$ in the reference population labelled naive $CD8^+$ T cells, which also included some double-negative T cells (Fig. 2f,g). The distribution placed the inferred program in lymphocyte populations with known effector functions. The individual inferred proteins showed distinct cell-type patterns, while the score summarized their coordinated effector program across lymphocyte populations.

We examined whether the three high-scoring populations also showed a coherent phenotype among the proteins actually measured in the same cells. These populations were fixed from the inferred-score summary before the measured-marker comparison. For each measured protein, abundances were averaged within biological sample and comparison group; the resulting ten high-minus-other differences were evaluated as paired observations, with multiple-testing correction across all 79 proteins. Mean paired CLR differences, divided by the across-cell standard deviation of each measured protein, were 1.101 for ITGB2, 1.611 for CD244, 1.500 for KLRG1 and 1.305 for NCAM1. All four were higher in the high-scoring group in every paired sample, with paired-test Benjamini–Hochberg-adjusted $P < 10^{-6}$ (Fig. 2h). This agreement supports a measured adhesion, receptor and effector-associated phenotype [27–29].

Natural killer-cell subsets provided a more resolved test of the inferred program. The comparison included 2,329 CD16-high cells and 131 CD56-high/CD16-low cells across ten paired samples. Using within-sample median inferred abundances, the mean standardized CD16-high minus CD56-high/CD16-low differences were 0.549 for PRF1, $-2.244$ for KLRD1 and $-1.142$ for EOMES. Each of these directions was consistent across all ten samples; GZMB and TBX21 did not show a corresponding significant paired difference (Fig. 2i). The heterogeneity among proteins is compatible with the distinct receptor and differentiation states of natural killer-cell subsets [30–32].

Together, the cell-type distribution, measured-marker agreement and subset structure provide complementary biological support for extending fragmented panels. They identify biologically plausible patterns that can inform subsequent targeted measurements, connecting profile extension to the guide stage.

#### Supplementary Note 4. Guiding informative measurements through protein-panel search

Finally, we asked whether protein relationships learned from fragmented profiles could guide the design of future measurements. Limited detection channels make panel composition consequential: a compact assay should retain information about proteins that cannot be measured simultaneously. We therefore selected 60 of 128 candidate proteins in Sparks2023 using frozen ProtCell reconstruction [33]. Candidate panels were scored by reconstruction of their omitted proteins, with lineage anchors retained. Comparison panels were obtained using QR-based selection and CytoVI [13, 46, 47].

To assess the selected measurements independently of the selector's own reconstruction model, we fitted ridge regression, nearest-neighbour regression and extremely randomized trees to each panel. Sparks2023 evaluation used the same 24,000 test cells and 39 targets absent from all three selected panels. Evaluating common targets prevents differences in which proteins remain unmeasured from determining the comparison. In ridge, nearest-neighbour and tree-decoder order, the ProtCell panel achieved mean protein-wise Spearman correlations of 0.616, 0.623 and 0.630, compared with 0.598, 0.605 and 0.622 for QR and 0.600, 0.606 and 0.623 for CytoVI. Corresponding median protein-wise RMSE values were 0.260, 0.253 and 0.244 for ProtCell, versus 0.276, 0.275 and 0.259 for QR and 0.271, 0.271 and 0.258 for CytoVI (Fig. 2j). Agreement across these independent decoders supports information retained by the selected proteins beyond the search model itself.

We then transferred the marker sets to Beneyto-Calabuig2023 [9], fitting decoders in Cohort A and evaluating 45,903 Cohort B cells over 80 common targets. The ProtCell panel achieved Spearman correlations of 0.390, 0.383 and 0.380 across the same decoder order, compared with 0.312, 0.318 and 0.316 for QR and 0.308, 0.309 and 0.307 for CytoVI. ProtCell median protein-wise RMSE ranged from 0.319 to 0.351, compared with 0.349–0.380 for QR and 0.353–0.375 for CytoVI. The transfer involved frozen marker choices, while the downstream decoders were fitted to the new dataset. Source panels all contained 60 markers, but native mapping retained 41, 39 and 40 markers for ProtCell, QR and CytoVI, respectively. Median protein-wise RMSE was calculated on the transformed expression values supplied to the decoders.

Matched random panels provided a further reference for the selected protein-wise summaries. The ProtCell panel exceeded the random-panel median Spearman correlation and had lower median RMSE for all three displayed decoders in both datasets. In Sparks2023, tree-decoder Spearman correlation was 0.6299 for ProtCell and 0.6297 at the random-panel median. Comparisons with alternative marker combinations further support the information retained by the selected measurements.

We separately tested whether the selected proteins retained information for acute myeloid leukaemia clone discrimination. For this analysis, each source panel was mapped to exactly 40 measured proteins using a fixed mapping and backfilling rule without leukaemia labels. A logistic regression classifier was fitted in Cohort A and evaluated in three held-out Cohort B patients. Classifier inputs consisted of these 40 measured proteins. Median patient-level AUROC and AUPRC were 0.809 and 0.968 for ProtCell, versus 0.478 and 0.786 for QR and 0.475 and 0.785 for CytoVI (Fig. 2k). Across 500 matched random panels, median AUROC and AUPRC were 0.739 and 0.855; ProtCell exceeded 70.0% of the random panels by AUROC and 96.8% by AUPRC. This measured-only result supports retention of classification-relevant information as well as completion information.

Together, these experiments connect the three stages of the framework: shared cell representations connect fragmented studies, contextual protein representations extend limited profiles, and the learned relationships help prioritize informative measurements. By retaining information for both profile completion and biological classification, panel search provides a basis for designing informative assays under limited detection capacity. Future studies can prospectively validate the selected protein combinations in new antibody panels and larger independent cohorts, and adapt the selection objectives to different cellular systems and experimental questions.

### Supplementary Tables

Supplementary Table 1: **Datasets retained for ProtCell pretraining.** Each row denotes one input file or panel, with identifiers and panel suffixes preserved. Links identify the originating study or data record. Cell and target counts describe the retained training inputs; cell counts were verified against the corresponding data shards. CyTOF inputs used arcsinh preprocessing (ARCSIN) and CITE-seq inputs used centered log-ratio preprocessing (CLR). Targets are harmonized input features, including combined or modification-specific identities.

| **Dataset ID** | **Study/source** | **Assay** | **Cells** | **Targets** | **Transform** |
|---|---|---|---:|---:|---|
| Dataset1 | Hartmann et al. (2019) | CyTOF | 2,567,549 | 32 | ARCSIN |
| Dataset2 | Hartmann et al. (2019) | CyTOF | 850,816 | 31 | ARCSIN |
| Dataset3 | Gonzalez et al. (2021) | CyTOF | 1,436,612 | 40 | ARCSIN |
| Dataset4 | Gonzalez et al. (2021) | CyTOF | 989,081 | 40 | ARCSIN |
| Dataset8 | Wu et al. (2021) | CyTOF | 2,799,097 | 35 | ARCSIN |
| Dataset9 | Suwandi et al. (2020) | CyTOF | 1,017,246 | 35 | ARCSIN |
| Dataset11 | Zheng et al. (2020) | CyTOF | 9,416,027 | 35 | ARCSIN |
| Dataset12 | Neidleman et al. (2020) | CyTOF | 447,760 | 29 | ARCSIN |
| Dataset13 | Ishiyama et al. (2022) | CyTOF | 25,672,460 | 36 | ARCSIN |
| Dataset14 | Khalsa et al. (2020) | CyTOF | 147,073 | 41 | ARCSIN |
| Dataset16 | Shi et al. (2022) | CyTOF | 1,769,076 | 34 | ARCSIN |
| Dataset17_1 | Hartmann et al. (2021) | CyTOF | 5,068,982 | 41 | ARCSIN |
| Dataset17_2 | Hartmann et al. (2021) | CyTOF | 1,472,530 | 41 | ARCSIN |
| Dataset17_3 | Hartmann et al. (2021) | CyTOF | 3,157,496 | 38 | ARCSIN |
| Dataset18 | Wei et al. (2022) | CyTOF | 1,678,856 | 34 | ARCSIN |
| Dataset21 | Ma et al. (2022) | CyTOF-Lec[a] | 2,227,443 | 32 | ARCSIN |
| Dataset22 | Stensland et al. (2022) | CyTOF | 6,306,390 | 37 | ARCSIN |
| Dataset23 | Stensland et al. (2022) | CyTOF | 4,017,312 | 33 | ARCSIN |
| Dataset24_1 | Ingelfinger et al. (2022) | CyTOF | 5,769,944 | 34 | ARCSIN |
| Dataset24_2 | Ingelfinger et al. (2022) | CyTOF | 8,830,398 | 33 | ARCSIN |
| Dataset26 | Miron et al. (2018) | CyTOF | 1,123,513 | 29 | ARCSIN |
| Dataset27_1 | Ivison et al. (2022) | CyTOF | 2,832,821 | 38 | ARCSIN |
| Dataset27_2 | Ivison et al. (2022) | CyTOF | 16,113,983 | 37 | ARCSIN |
| Dataset28 | Han et al. (2019) | CyTOF | 19,302,445 | 33 | ARCSIN |
| Dataset30 | Ma et al. (2021) | CyTOF | 7,264,703 | 34 | ARCSIN |
| Dataset31 | Georg et al. (2022) | CyTOF | 23,247,998 | 30 | ARCSIN |
| Dataset36 | Ma et al. (2020) | CyTOF | 7,643,411 | 37 | ARCSIN |
| Dataset37_1 | Sponaugle et al. (2022) | CyTOF | 26,797,163 | 33 | ARCSIN |
| Dataset37_2 | Sponaugle et al. (2022) | CyTOF | 8,867,677 | 33 | ARCSIN |
| Dataset39 | Neidleman et al. (2020) | CyTOF | 4,096,542 | 36 | ARCSIN |
| Dataset42 | Roan (2022) | CyTOF | 751,516 | 37 | ARCSIN |
| Dataset43 | Neidleman et al. (2021) | CyTOF | 12,721,676 | 35 | ARCSIN |
| Dataset44 | Melenhorst et al. (2022) | CyTOF | 45,117 | 37 | ARCSIN |
| Dataset45 | George et al. (2022) | CyTOF | 8,176,401 | 37 | ARCSIN |
| Dataset47 | Xie et al. (2021) | CyTOF | 5,473,136 | 36 | ARCSIN |
| Dataset49_1 | Vallve-Juanico et al. (2022) | CyTOF | 3,882,837 | 40 | ARCSIN |
| Dataset49_2 | Vallve-Juanico et al. (2022) | CyTOF | 8,793,826 | 40 | ARCSIN |
| Dataset52_1 | Li et al. (2020) | CyTOF | 3,393,688 | 35 | ARCSIN |

Continued on the next page

**Supplementary Table 1 continued**

| Dataset ID | Study/source | Assay | Cells | Targets | Transform |
|---|---|---|---|---|---|
| Dataset52_2 | Li et al. (2020) | CyTOF | 47,294 | 35 | ARCSIN |
| Dataset53_1 | Kim et al. (2022) | CyTOF | 25,567 | 40 | ARCSIN |
| Dataset53_2 | Kim et al. (2022) | CyTOF | 25,912 | 41 | ARCSIN |
| Dataset53_3 | Kim et al. (2022) | CyTOF | 25,833 | 40 | ARCSIN |
| Dataset53_4 | Kim et al. (2022) | CyTOF | 26,010 | 41 | ARCSIN |
| Dataset53_5 | Kim et al. (2022) | CyTOF | 24,504 | 39 | ARCSIN |
| Dataset53_6 | Kim et al. (2022) | CyTOF | 26,133 | 40 | ARCSIN |
| Dataset53_7 | Kim et al. (2022) | CyTOF | 21,813 | 38 | ARCSIN |
| Dataset53_8 | Kim et al. (2022) | CyTOF | 20,009 | 37 | ARCSIN |
| Dataset53_9 | Kim et al. (2022) | CyTOF | 19,772 | 40 | ARCSIN |
| Dataset53_10 | Kim et al. (2022) | CyTOF | 23,170 | 38 | ARCSIN |
| Dataset53_11 | Kim et al. (2022) | CyTOF | 18,758 | 39 | ARCSIN |
| Dataset53_12 | Kim et al. (2022) | CyTOF | 21,158 | 31 | ARCSIN |
| Dataset53_13 | Kim et al. (2022) | CyTOF | 16,466 | 38 | ARCSIN |
| Dataset53_14 | Kim et al. (2022) | CyTOF | 19,559 | 37 | ARCSIN |
| Dataset53_15 | Kim et al. (2022) | CyTOF | 19,125 | 30 | ARCSIN |
| Dataset54_2 | Roussel et al. (2021) | CyTOF | 13,746,884 | 34 | ARCSIN |
| Dataset55_1 | Chevrier et al. (2021) | CyTOF | 2,241,053 | 40 | ARCSIN |
| Dataset55_2 | Chevrier et al. (2021) | CyTOF | 417,403 | 40 | ARCSIN |
| Dataset55_3 | Chevrier et al. (2021) | CyTOF | 218,996 | 40 | ARCSIN |
| Dataset56 | Rodriguez et al. (2020) | CyTOF | 6,632,021 | 47 | ARCSIN |
| Dataset60_1 | Friebel et al. (2020) | CyTOF | 16,754,114 | 37 | ARCSIN |
| Dataset61 | Wang et al. (2022) | CyTOF | 1,770,339 | 33 | ARCSIN |
| Dataset63 | Petrilli et al. (2022) | CyTOF | 342,919 | 17 | ARCSIN |
| Dataset65 | Michlmayr et al. (2018) | CyTOF | 12,475,661 | 36 | ARCSIN |
| Dataset76_1 | Guilliams et al. (2022) | CITE-seq | 73,580 | 179 | CLR |
| Dataset78_1.2 | Maier et al. (2020) | CITE-seq | 588 | 57 | CLR |
| Dataset78_1.3 | Maier et al. (2020) | CITE-seq | 1,254 | 79 | CLR |
| Dataset80_1 | Liu et al. (2021) | CITE-seq | 246,964 | 183 | CLR |
| Dataset80_2 | Liu et al. (2021) | CITE-seq | 125,117 | 183 | CLR |
| Dataset81 | Pei et al. (2020) | CITE-seq | 4,426 | 19 | CLR |
| Dataset82_1 | Bai et al. (2022) | CITE-seq | 23,287 | 16 | CLR |
| Dataset82_2 | Bai et al. (2022) | CITE-seq | 31,105 | 16 | CLR |
| Dataset82_3 | Bai et al. (2022) | CITE-seq | 30,484 | 16 | CLR |
| Dataset82_4 | Bai et al. (2022) | CITE-seq | 16,450 | 16 | CLR |
| Dataset88 | Witkowski et al. (2020) | CITE-seq | 44,041 | 77 | CLR |
| Dataset91 | Nathan et al. (2021) | CITE-seq | 500,089 | 30 | CLR |
| Dataset92 | Kumar et al. (2022) | CITE-seq | 85,000 | 12 | CLR |
| Dataset93 | Lasry et al. (2023) | CITE-seq | 211,442 | 259 | CLR |
| Dataset94 | Unterman et al. (2022) | CITE-seq | 38,652 | 180 | CLR |
| Dataset95 | Vanuytsel et al. (2022) | CITE-seq | 26,407 | 16 | CLR |
| Dataset100 | Kaufmann et al. (2021) | CITE-seq | 355,433 | 37 | CLR |
| Dataset117 | Paschalidis et al. (2025) | CyTOF | 1,666,570 | 33 | ARCSIN |
| Dataset118 | Wegmann (2024) | CyTOF | 3,075,527 | 40 | ARCSIN |
| Dataset120 | Roan (2023) | CyTOF | 4,558,842 | 34 | ARCSIN |
| Dataset123_1 | Wing (2024), Zenodo | CyTOF | 3,112,266 | 36 | ARCSIN |
| Dataset123_2 | Wing (2024), Zenodo | CyTOF | 24,945,646 | 34 | ARCSIN |

Continued on the next page

**Supplementary Table 1 continued**

| Dataset ID | Study/source | Assay | Cells | Targets | Transform |
|---|---|---|---|---|---|
| Dataset123_3 | Wing (2024), Zenodo | CyTOF | 42,605,735 | 39 | ARCSIN |
| **Total: 86 retained files/panels** | | | **382,929,979** | – | – |

**Counting and preprocessing.** The training inputs comprise 69 CyTOF files with 381,115,660 cell records and 17 CITE-seq files with 1,814,319 cell records, covering 616 distinct target features. Only the preprocessing version matching each technology contributes to these totals. Counts refer to input cell records rather than independent donors or studies; panels from one publication remain separate rows. Per-file target counts describe retained input features, rather than original antibody-panel sizes or the full vocabulary of 646 tokens, which includes two special tokens. Full source citations and the correspondence between training inputs and source records are provided in the accompanying Supplementary Table 1 workbook.

[a] Dataset21 is annotated CyTOF-Lec in the source worksheet and grouped under CyTOF in the training metadata.

### Supplementary references


[7] Johanna Wagner et al. A single-cell atlas of the tumor and immune ecosystem of human breast cancer. *Cell*, 177:1330–1345.e18, 2019.

[8] Yuri Kotliarov et al. Broad immune activation underlies shared set point signatures for vaccine responsiveness in healthy individuals and disease activity in patients with lupus. *Nature Medicine*, 26:618–629, 2020.

[9] Sergi Beneyto-Calabuig et al. Clonally resolved single-cell multi-omics identifies routes of cellular differentiation in acute myeloid leukemia. *Cell Stem Cell*, 30:706–721.e8, 2023.

[12] Sofie Van Gassen et al. Flowsom: using self-organizing maps for visualization and interpretation of cytometry data. *Cytometry Part A*, 87:636–645, 2015.

[13] Florian Ingelfinger, Nathan Levy, Can Ergen, et al. CytoVI: deep generative modeling of antibody-based single cell data. *Nature Methods*, 23:2054–2065, 2026.

[14] Alexandre Segers et al. omicsgmf: a multi-tool for dimensionality reduction, batch correction and imputation in bulk- and single-cell proteomics. *Nature Communications*, 17:6650, 2026.

[17] Keith Sacco et al. Immunopathological signatures in multisystem inflammatory syndrome in children and pediatric covid-19. *Nature Medicine*, 28:1050–1062, 2022.

[18] Federica Sallusto, Danielle Lenig, Reinhold Förster, Martin Lipp, and Antonio Lanzavecchia. Two subsets of memory t lymphocytes with distinct homing potentials and effector functions. *Nature*, 401:708–712, 1999.

[19] Luca Gattinoni et al. A human memory t cell subset with stem cell-like properties. *Nature Medicine*, 17:1290–1297, 2011.

[20] N. K. Damle and L. V. Doyle. Ability of human T lymphocytes to adhere to vascular endothelial cells and to augment endothelial permeability to macromolecules is linked to their state of post-thymic maturation. *The Journal of Immunology*, 144(4):1233–1240, 1990.

[21] Laura S. Peterson et al. Single-cell analysis of the neonatal immune system across the gestational age continuum. *Frontiers in Immunology*, 12:714090, 2021.

[22] John H. Russell and Timothy J. Ley. Lymphocyte-mediated cytotoxicity. *Annual Review of Immunology*, 20:323–370, 2002.

[23] Ilia Voskoboinik, James C. Whisstock, and Joseph A. Trapani. Perforin and granzymes: function, dysfunction and human pathology. *Nature Reviews Immunology*, 15:388–400, 2015.

[24] Eric Vivier, Elena Tomasello, Myriam Baratin, Thierry Walzer, and Sophie Ugolini. Functions of natural killer cells. *Nature Immunology*, 9:503–510, 2008.

[25] Susan M. Kaech and Weiguo Cui. Transcriptional control of effector and memory cd8+ t cell differentiation. *Nature Reviews Immunology*, 12:749–761, 2012.

[26] Scott M. Gordon et al. The transcription factors t-bet and eomes control key checkpoints of natural killer cell maturation. *Immunity*, 36:55–67, 2012.

[27] Jordan S. Orange. Formation and function of the lytic NK-cell immunological synapse. *Nature Reviews Immunology*, 8(9):713–725, 2008.

[28] Nikhil S. Joshi et al. Inflammation directs memory precursor and short-lived effector CD8+ T cell fates via the graded expression of T-bet transcription factor. *Immunity*, 27(2):281–295, 2007.

[29] Michael A. Caligiuri. Human natural killer cells. *Blood*, 112(3):461–469, 2008.

[30] Jianhua Yu et al. CD94 surface density identifies a functional intermediary between the CD56bright and CD56dim human NK-cell subsets. *Blood*, 115(2):274–281, 2010.

[31] James J. Knox, Gabriela L. Cosma, Michael R. Betts, and Laura M. McLane. Characterization of T-bet and Eomes in peripheral human immune cells. *Frontiers in Immunology*, 5:217, 2014.

[32] Julia A. Wagner et al. CD56bright NK cells exhibit potent antitumor responses following IL-15 priming. *The Journal of Clinical Investigation*, 127(11):4042–4058, 2017.

[33] Rachel Sparks et al. Influenza vaccination reveals sex dimorphic imprints of prior mild COVID-19. *Nature*, 614(7949):752–761, 2023.

[37] Dvir Aran et al. Reference-based analysis of lung single-cell sequencing reveals a transitional profibrotic macrophage. *Nature Immunology*, 20:163–172, 2019.

[38] Tim Stuart et al. Comprehensive integration of single-cell data. *Cell*, 177:1888–1902.e21, 2019.

[46] Peter Businger and Gene H. Golub. Linear least squares solutions by Householder transformations. *Numerische Mathematik*, 7:269–276, 1965.

[47] Krithika Manohar, Bingni W. Brunton, J. Nathan Kutz, and Steven L. Brunton. Data-driven sparse sensor placement for reconstruction: Demonstrating the benefits of exploiting known patterns. *IEEE Control Systems Magazine*, 38(3):63–86, 2018.
